# The global distribution of *Bacillus anthracis* and associated anthrax risk to humans, livestock, and wildlife

**DOI:** 10.1101/394023

**Authors:** Colin J. Carlson, Ian T. Kracalik, Noam Ross, Kathleen Alexander, Martin E. Hugh-Jones, Mark Fegan, Brett Elkin, Tasha Epp, Todd K. Shury, Mehriban Bagirova, Wayne M. Getz, Jason K. Blackburn

**Affiliations:** National Socio-Environmental Synthesis Center, University of Maryland, Annapolis, Maryland 21401, USA. Department of Biology, Georgetown University, Washington, D.C. 20057, USA.; Spatial Epidemiology & Ecology Research Lab, Department of Geography, University of Florida, Gainesville, FL, USA Emerging Pathogens Institute, University of Florida, Gainesville, FL, USA.; EcoHealth Alliance, 460 W 34th St FL 17, New York, NY 10001, USA.; Department of Fish and Wildlife Conservation, Virginia Tech, Cheatham Hall, Room 101, Blacksburg, VA, USA.; School of the Coast and Environment, Louisiana State University, Baton Rouge, LA, USA.; AgriBio, Centre for Agribiosciences, Biosciences Research, Department of Economic Development, Jobs, Transport and Resources, Bundoora Victoria, Australia.; Department of Environment and Natural Resources, Government of the Northwest Territories, 5102-50th Ave., Yellowknife, Northwest Territories X1A 3S8, Canada.; Department of Large Animal Clinical Sciences, Western College of Veterinary Medicine, University of Saskatchewan, Saskatoon, Canada.; Parks Canada Agency, 52 Campus Dr., Saskatoon, Saskatchewan S7N 5B4, Canada.; Scientific Research Veterinary Institute, Baku, Azerbaijan.; Department of Environmental Science, Policy, and Management, University of California, Berkeley, Berkeley, California, USA.

## Abstract

*Bacillus anthracis* is a spore-forming, Gram-positive bacterium responsible for anthrax, an acute and commonly lethal infection that most significantly affects grazing livestock, wild ungulates and other herbivorous mammals, but also poses a serious threat to human health^1, 2^. The geographic extent of *B. anthracis* endemism is still poorly understood, despite multi-decade research on anthrax epizootic and epidemic dynamics around the world^3, 4^. Several biogeographic studies have focused on modeling environmental suitability for anthrax at local or national scales^5–9^, but many countries have limited or inadequate surveillance systems, even within known endemic regions. Here we compile an extensive global occurrence dataset for *B. anthracis*, drawing on confirmed human, livestock, and wildlife anthrax outbreaks. With these records, we use boosted regression trees^10, 11^ to produce the first map of the global distribution of *B. anthracis* as a proxy for anthrax risk. Variable contributions to the model support pre-existing hypotheses that environmental suitability for *B. anthracis* depends most strongly on soil characteristics such as pH that affect spore persistence, and the extent of seasonal fluctuations in vegetation, which plays a key role in transmission for herbivores^12, 13^. We apply the global model to estimate that 1.83 billion people (95% credible interval: 0.59—4.16 billion) live within regions of anthrax risk, but most of that population faces little occupational exposure to anthrax. More informatively, a global total of 63.8 million rural poor livestock keepers (95% CI: 17.5—168.6 million) and 1.1 billion livestock (95% CI: 0.4—2.3 billion) live within vulnerable regions. Human risk is concentrated in rural areas, and human and livestock vulnerability are both concentrated in rainfed systems throughout arid and temperate land across Eurasia, Africa, and North America. We conclude by mapping where anthrax risk overlaps with vulnerable wild ungulate populations, and therefore could disrupt sensitive conservation efforts for species like bison, pronghorn, and saiga that coincide with anthrax-prone, mixed-agricultural landscapes. Anthrax is a zoonotic disease caused by the Gram-positive bacterium *Bacillus anthracis*, a generalist soil-transmitted pathogen found on every inhabited continent^14^, and several islands including Haiti and parts of the Philippines and Indonesia. Worldwide, an estimated 20,000 to 100,000 cases of anthrax occur annually, mostly in poor rural areas^15^. In clinical presentations of anthrax, case fatality rates are a function of exposure pathway. Respiratory exposure from spore inhalation is important the context of bioterrorism, but is highly uncommon, and accounts for a negligible fraction of the global burden of anthrax cases. Cutaneous exposure to *B. anthracis* accounts for the majority of human cases worldwide, and typically presents with low mortality; gastrointestinal exposure accounts for the remainder and presents with intermediate to high fatality rates. Cutaneous and gastrointestinal cases of anthrax are most commonly caused by handling and slaughtering infected livestock, or butchering and eating contaminated meat; untreated gastrointestinal cases likely account for most human mortality from anthrax.^14–16^

---

Human mortality from anthrax is driven by ecological dynamics at the wildlife-livestock interface^17^. In nature, the enzootic cycle of anthrax is characterized by a combination of long-term spore persistence in soil, and an obligate-lethal transmission route, primary in herbivorous mammals^1, 2, 14^. Both wild herbivores and livestock are gastrointestinally exposed to *B. anthracis* spores from soil while grazing, become infected, and usually return spores to the soil when they die and decompose^13^. Because domesticated and wild herbivores frequently share grazing grounds, wildlife epizootics can lead to downstream infections in livestock and humans. In some regions, anthrax is hyperendemic, and cases follow regular seasonal trends; in other regions, the disease re-emerges in major epidemics after years or decades without a single case^1^. Environmental persistence facilitates these unusual dynamics; under optimal conditions, *B. anthracis* spores are able to persist in the soil for long periods (i.e. decades). Alkaline, calcium-rich soils are believed to facilitate sporulation, and therefore drive landscape-level patterns of persistence; these patterns are usually hypothesized to drive the distribution of *B. anthracis* up to the continental scale^2, 14^.

Global variation in anthrax endemism and outbreak intensity has been previously characterized at extremely coarse scales^3, 18^, but anthrax is a neglected disease, and its global distribution is still poorly characterized. In total, roughly a dozen studies have used ecological niche models to develop regional (usually national-level) maps of suitability for *B. anthracis* (see **Table S1**). These regional mapping efforts are a critical part of the public health planning process^4^, but are primarily conducted in isolation, and the results of these studies have yet to be consolidated and synthesized. Furthermore, cross-validation of regional models has only been recently attempted^19^, and indicates either limitations in model transferability, or possible genetic or ecological differences underlying distributional patterns of different regions; either way, this highlights the limitations preventing regional models from being scaled up to a global estimate. Moreover, the distribution of anthrax has yet to be modelled in several broad regions where it is nevertheless pervasive, especially Western Europe, the Middle East, and South America. Cryptic persistence of *B. anthracis* spores in the soil makes mapping efforts especially challenging, as suitable and endemic regions could go years or potentially decades without a recorded outbreak.

This study consolidates clinical and ecological research on enzootic and epidemic anthrax reports, compiling the largest global database of anthrax occurrences on record to map the global suitability for *Bacillus anthracis* persistence. A total of 5,108 records were compiled describing the global distribution of anthrax across 70 countries (**Figure 1**). Here we used a subset of 2,310 of these data points to describe the global distribution and eco-epidemiology of *Bacillus anthracis*, exploring the relationship of anthrax outbreaks to environmental factors including soil characteristics and climate, via boosted regression trees (BRTs) as a tool for species distribution modelling. These maps provide a proxy for anthrax risk. We apply this global anthrax model to provide a first estimate of the global human and livestock populations at risk from anthrax. We compare the distribution of anthrax to that of critically threatened wildlife populations, and identify areas where additional or new surveillance is needed to anticipate and prevent rare, but likely catastrophic, threats to wildlife conservation efforts.

**Figure 1.**
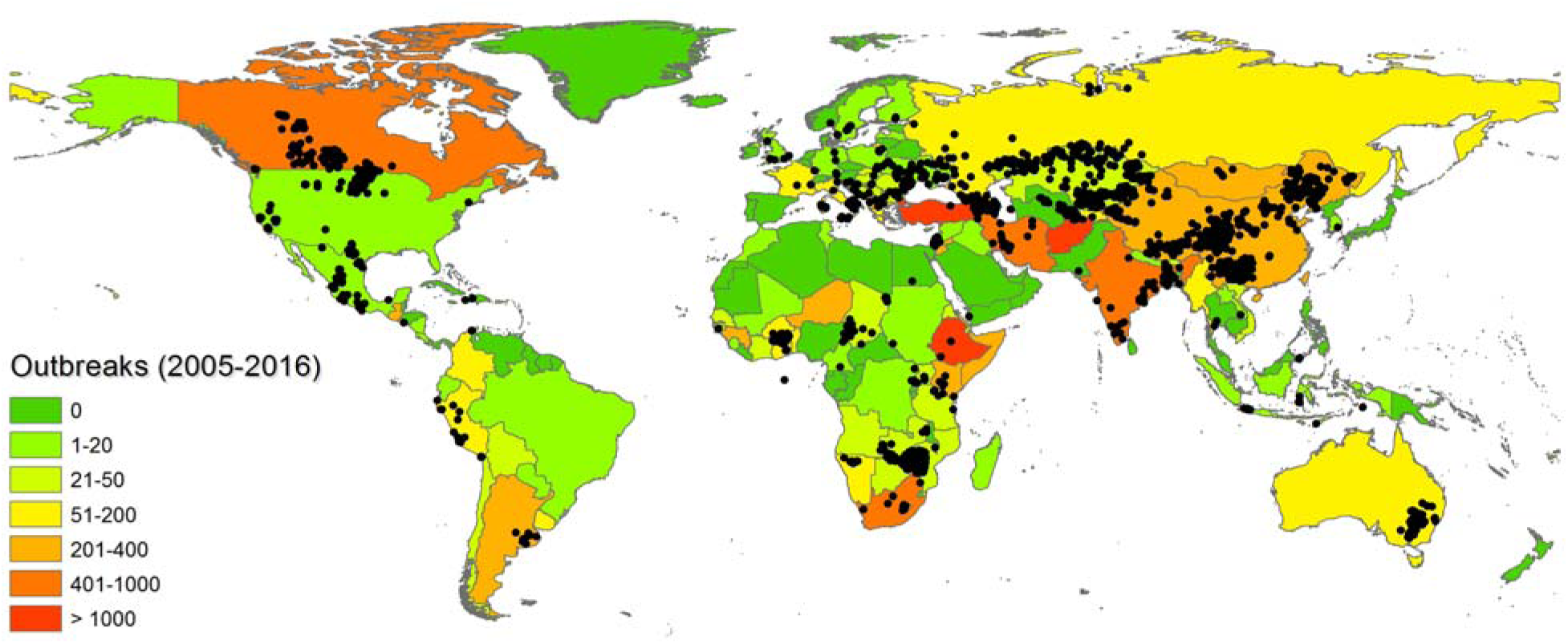
Global database of anthrax occurrences (points), versus outbreaks of anthrax by country (Jan. 2005-Aug. 2016; data digitized from United Nations Food & Agriculture Organization report).

Our global ensemble distribution model (**Figure 2**) performed very well on validation data (mean AUC = 0.9244), and regionally matches the well-established distribution of anthrax in China^20^, Kazakhstan^21^, North America^5^, and Australia^8^, suggesting that it appropriately captures the global range of *Bacillus anthracis*. These four regions, along with a band of suitability in sub-Saharan Africa around roughly 15° S, represent the most geographically-expansive zones of anthrax endemicity, and hotspots of human vulnerability. However, our study also shows that perhaps the majority of the European continent, and a substantial part of the Anatolian peninsula and surrounding region, are also highly suitable for *B. anthracis.* In some cases, high risk areas match hyperendemic areas, such as Turkey and South Africa; but in other cases (e.g., in Ethiopia), predicted areas of suitability were more limited than expected in countries with a high anthrax burden. This may reflect sampling limitations in these areas, but also likely reflects regional variation in anthrax control. Where food safety practices prevent exposure and livestock vaccinations are affordable, high anthrax suitability may not translate into high case burdens (supported by the vaccination data in the **Supporting Information**); conversely, anthrax morbidity and mortality are usually exacerbated by limited local knowledge about anthrax, limited access to healthcare, and conflicting pressures like food insecurity that force local populations to handle and eat contaminated meat.

**Figure 2.**
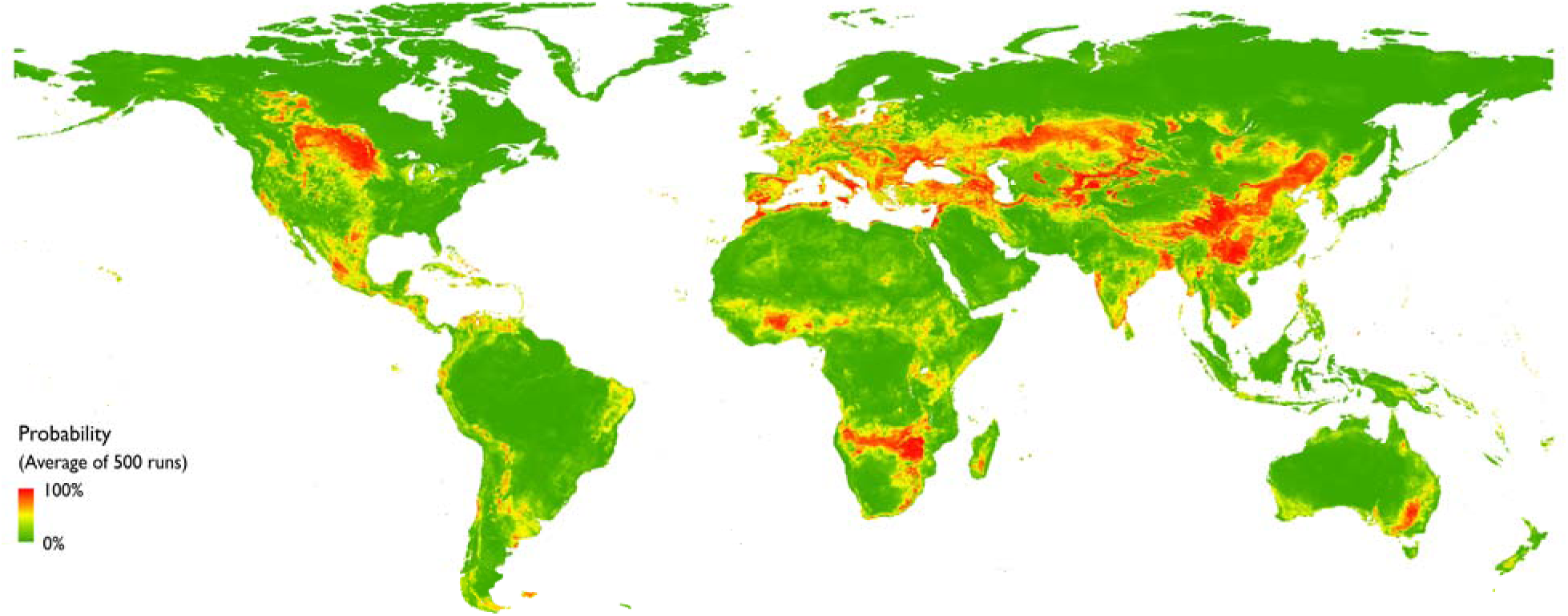
Global environmental suitability (probability of occurrence) for *Bacillus anthracis* modelled by an ensemble of 500 boosted regression trees.

**Figure 3.**
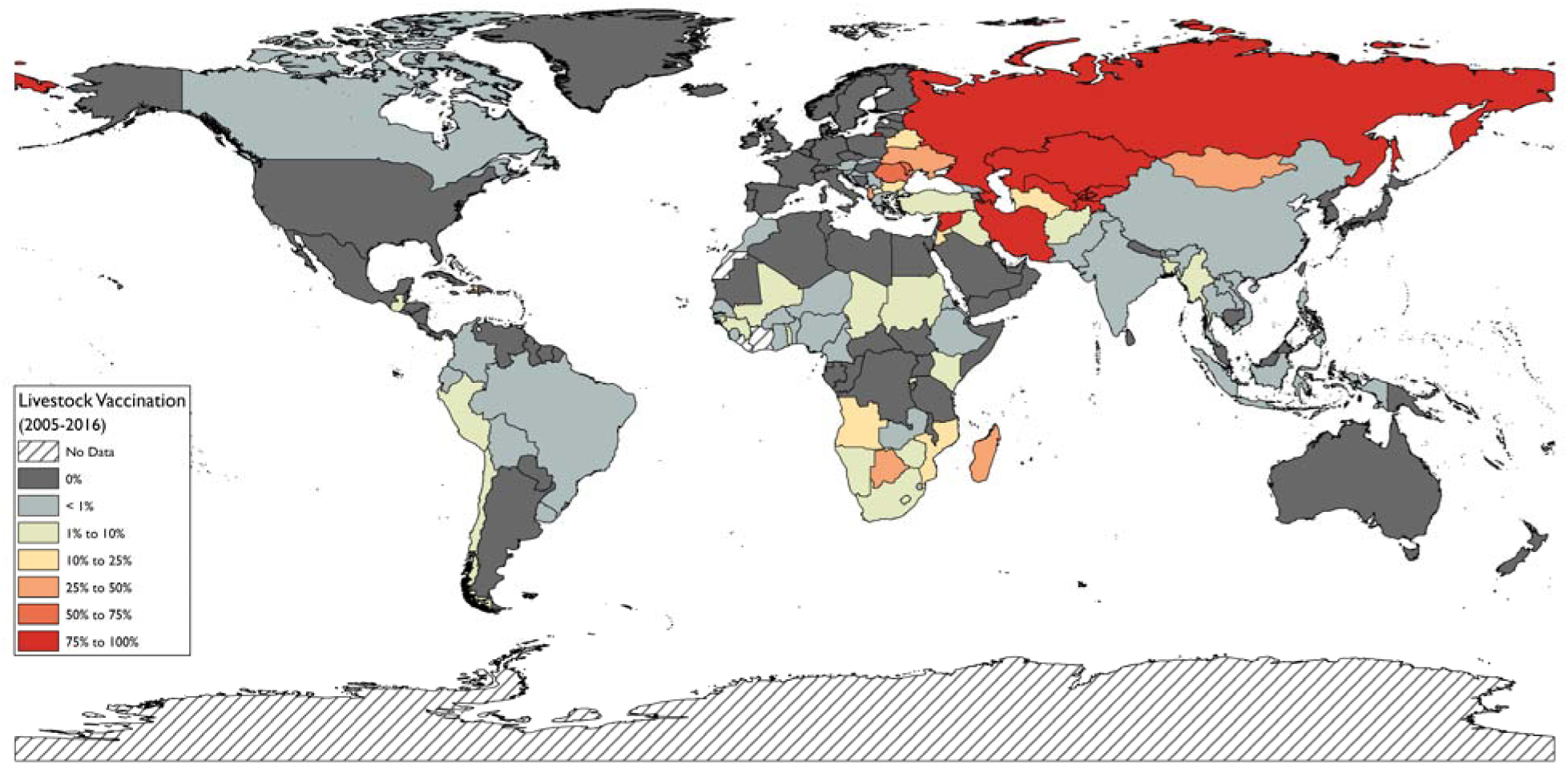
Average vaccination rates for all livestock (cattle, sheep, goats, buffalo, and pigs) reported to WAHIS over the interval 2005-2016, as a function of doses administered and reported livestock populations at risk, excluding years with no reported vaccination.

Globally, we find an estimated 1.8 billion people live within anthrax-suitable areas, the vast majority of whom live in rural areas in Africa, Europe and Asia (**Table 1**). However, most of that population probably has no occupational exposure to infected animals, and direct exposure from the soil has rarely been reported for human cases; in those few reported cases patients had compromised immune systems and unknown or unusual exposure^22^. For a more informative perspective on risk, we estimate that a total of 63.8 million rural poor livestock keepers live in anthrax-affected areas (95% credible interval: 17.5—168.6 million; **Table 1**), again primarily in Africa and Eurasia. Globally, we found that areas of anthrax risk contain 1.1 billion livestock (95% CI: 404 million—2.3 billion; **Table 2**), including 320 million sheep (95% CI: 138—622 million), 294.9 million pigs (95% CI: 103—583 million), 268.1 million cattle (95% CI: 87.4—639 million), 211.2 million goats (95% CI: 74.8—453 million), and 0.6 million buffalo (95% CI: 0.16—1.6 million). Although arid and semi-arid ecosystems were a significant source of vulnerable livestock across production systems (especially for cattle), the single most significant across all four groups was rainfed mixed crop/livestock systems in temperate-highland ecosystems, due to the disproportionate contribution from East Asia, especially China (Table 1).

**Table 1.**
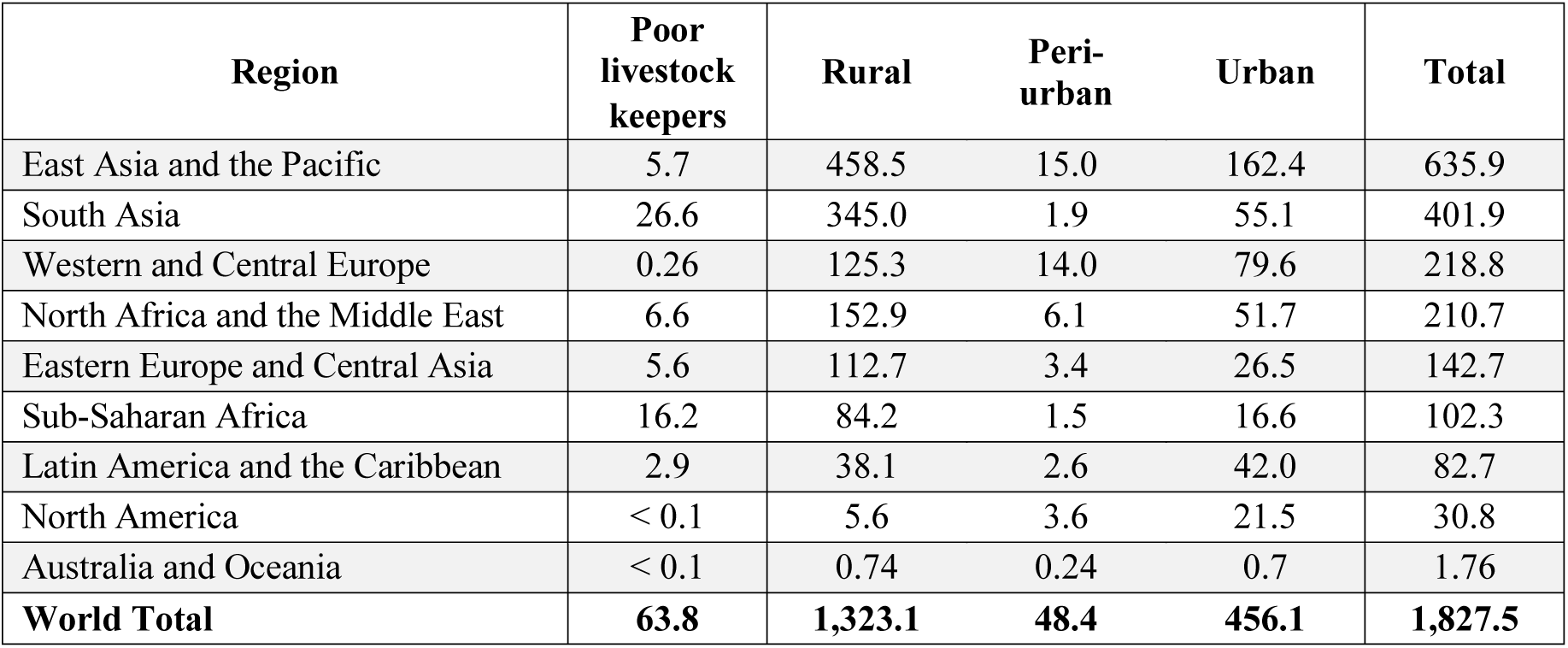
Population at risk (in millions) by region, land use, and occupational exposure.

**Table 2.** Estimated global livestock at risk (by millions), by species & region.

| **Region** | **Cattle** | **Pigs** | **Goats** | **Sheep** | **Buffalo** | **Total** |
| --- | --- | --- | --- | --- | --- | --- |
| East Asia and the Pacific | 63.2 | 190.9 | 79.9 | 108.1 | 0.24 | 442.4 |
| South Asia | 61.6 | 1.8 | 72.5 | 18.7 | 0.33 | 154.8 |
| Western and Central Europe | 22.2 | 60.9 | 7.5 | 42.2 | < 0.1 | 132.8 |
| North Africa and the Middle East | 15.8 | 0.2 | 13.4 | 65.2 | < 0.1 | 94.6 |
| Eastern Europe and Central Asia | 26.6 | 12.0 | 8.9 | 40.4 | < 0.1 | 87.8 |
| Sub-Saharan Africa | 30.5 | 5.3 | 22.4 | 14.5 | 0 | 72.8 |
| Latin America and the Caribbean | 21.9 | 8.0 | 5.7 | 8.1 | 0 | 43.7 |
| North America | 23.0 | 15.2 | 0.45 | 0.29 | 0 | 39.0 |
| Australia and Oceania | 3.1 | 0.61 | 0.47 | 22.7 | 0 | 26.9 |
| **World Total** | **268.1** | **294.9** | **211.2** | **320.1** | **0.58** | **1,094.9** |

Most livestock at risk of anthrax are unvaccinated in any given year. Per reported data, an average of 212.8 million live attenuated vaccines for anthrax are manufactured every year (2005-2016) outside the United States (which reports no data but is also a major manufacturer); on average, 198.2 million doses are used for livestock every year. Compared to the 1.1 billion livestock at risk, vaccine coverage is patchy and regionally-variable; roughly 90% of cattle, sheep, and goats are annually vaccinated in Eastern Europe and Central Asia, due to a strong legacy of mass vaccination campaigns in the former Soviet Union. On the other hand, vaccination rates are alarmingly low in sub-Saharan Africa (0-6%), East Asia (0-5%) and South Asia (< 1%), where more than half of the livestock at risk and 48.5 million rural poor livestock keepers are located. In these regions, livestock vaccination is commonly used reactively after a major outbreak, rather than as a preventative measure^5, 23^; improving proactive vaccination in under-vaccinated, hyperendemic countries (in particular Afghanistan, Bangladesh, Ethiopia, South Africa, Turkey, and Zimbabawe) could help bring anthrax outbreaks under control^24^. Vaccination may also be less effective than usual for the 31 million livestock and 4.6 million poor livestock keepers in West Africa, where an endemic lineage of *B. anthracis* shares an anthrose-deficiency mutation that has been hypothesized to lead to a vaccine escape^25^. Education campaigns may be more cost-effective than mass vaccination, which is both cost-prohibitive and logistically-challenging in inaccessible rural areas. However, livestock keepers may continue to sell contaminated meat to recoup financial losses (which also contributes to underreporting); this has increased cases urban settings^26^. In cases of extreme food insecurity, poor populations may eat anthrax-infected meat despite understanding the risks.

Although risk is most commonly measured at the human-agriculture interface, anthrax also has a major ecological impact; while *B. anthracis* is a stable part of some savannah ecosystems, epizootics in other regions can have catastrophic impacts on wildlife populations^14, 27^. We note that several ungulate species could probably benefit from improved epizootic surveillance, given range overlap with anthrax and limited coverage by protected areas, which are a foundation of anthrax surveillance and control for most wildlife (**Table 3**).^28^ Saiga antelope (*Saiga tatarica*) in particular have a significant overlap with anthrax outside of protected parts of their range, and the recent mass die-off of a third of the entire population of saiga in three weeks highlights the vulnerability of threatened ungulates to sudden, disease-induced population crashes. The anthrax vaccine may be used by conservationists in special cases (e.g. with cheetahs and rhinoceros^29^), but the lack of an oral anthrax vaccine makes mass vaccination more impractical for wildlife than for livestock, making surveillance all the more important.

**Table 3.** Overlap between wildlife species of concern and the global distribution of anthrax, including overlap with the protected areas database.

| Wildlife Species | Geographic Overlap | Anthrax suitable area overlap with protected areas | IUCN Red List Status |
| --- | --- | --- | --- |
| Bison (*Bison bison*) | 32.2% | 64.6% | NT |
| Pronghorn (*Antilocapra americana*) | 26.6% | 3.9% | LC |
| Roan (*Hippotragus equinus*) | 23.9% | 30.5% | LC |
| Saiga (*Saiga tatarica*) | 17.5% | 2.8% | CE (A2acd) |
| Moose (*Alces alces*) | 6.0% | 9.9% | LC |
| Reindeer (*Rangifer tarandus*) | 1.6% | 14.9% | LC |
| Yak (*Bos mutus*) | 0.7% | 0% | VU (C10) |

Our study has proposed the first global map of *B. anthracis* suitability as a proxy for anthrax risk, and while this is a major step forward, several important directions remain to make these models actionable for public health practitioners. Although some estimates have been proposed for the annual global burden of anthrax, these estimates range by several orders of magnitude. Most regional assessments, especially in rural Africa, agree that anthrax cases are severely underreported despite mandatory reporting. Similar studies to ours have used suitability maps to extrapolate global case burden for diseases like dengue fever or melioidosis (*Burkholderia pseudomallei*)^11, 30^; however, this approach seems inadequate for anthrax, given that human incidence is just as strongly determined by anthrax dynamics in wildlife, local agricultural intensity, knowledge about anthrax transmission, access to healthcare and vaccination, and complicating factors like food insecurity. At national and local levels, One Health approaches to surveillance have had promising results, but a more globally-coordinated network among these programs might help address some of the major data gaps.

Our work also sets a foundation for investigating how climate change will impact the distribution and burden of anthrax. Published work suggests anthrax suitability may decrease in parts of Kazakhstan and the southern United States in a changing climate, but other work anticipates warming-driven emergence at higher latitudes^31, 32^. Our study includes recent records from the Yamalo-Nemets area of Russia, where outbreaks in reindeer have led to massive economic losses and threaten the livelihood of traditional pastoralists. However, our model made limited predictions of suitability in the sub-Arctic in current climates. Even though anthrax cases are regularly reported throughout Sweden, northern Russia, and other cold, high-latitude countries, high-latitude outbreaks are proportionally under-represented in our database (and are often poorly documented). Persistence at higher latitudes may also be better predicted by a slightly different set of climatic constraints on persistence. A recently-published model trained on high-latitude cases in the Northern Hemisphere seems to under-predict known areas of endemism in warmer climates, possibly supporting this explanation^32^. Feedback between local modelling efforts and updated global consensus mapping will improve our overall understanding of what drives different anthrax dynamics, and the likely impact of climate change.

Finally, we observe increasing interest, by microbiologists and ecologists alike, in the closely related “anthrax-like” *Bacillus cereus* biovar. *anthracis.* Whereas *Bacillus cereus* is a typically non-pathogenic soil bacterium, the pathogenic *B. cereus* biovar. *anthracis* (*Bcbva*) carries variants of the pXO1 and pXO2 plasmids that allow capsule production, which are both required for full virulence in *B. anthracis.* A recent study in Taï National Park in Cote D’Ivôire showed that *Bcbva* accounted for 40% of wildlife mortality in a 26-year survey, and could potentially drive the local extinction of chimpanzees in the Taï Forest within the next century.^33^ The geographic distribution of *Bcbva* is still unknown, and it is plausible that different climatic and environmental factors determine the spatial patterns of its transmission; though improved diagnostics will be necessary to differentiate the role of the two pathogens in anthrax infections beyond Taï National Park. Mapping *Bcbva* across West Africa may be an important next step for measuring the threat of anthrax and anthrax-like disease to wildlife conservation (and, potentially, to human health down the road).

## Methods

### Occurrence & Pseudoabsence Data

We assembled a global occurrence database for *Bacillus anthracis* out of a combination of (i) outbreak data collected in the field by the authors or their extended team of collaborators, (ii) national passive surveillance and reporting infrastructures, (iii) online records from ProMed Mail, and (iv) georeferenced records or (v) digitized maps from peer-reviewed publications documenting anthrax outbreaks. Our final database of 5,018 unique records spanning 70 countries and more than a century (1914 – 2018), thinned to 2,310 distinct localities after removing uncertain sightings and thinning to a single point per unique 10 arcminute (∼20 km at the equator) cell, to correct for sampling bias^34^. For background environmental data, a total dataset of 10,000 pseudoabsences were randomly generated from countries where anthrax occurrence records were collected, an upper sample commonly used for similar disease distribution mapping studies^35^. Of these 10,000, an equal 1:1 sample was randomly selected to match the presence points in submodels, as is recommended for boosted regression tree models^36^.

### Environmental Predictors

Global layers of environmental predictors were selected based on successful variables used in previously published anthrax distribution modelling studies at regional scales^5, 21, 37^, as well as from other distribution modelling studies on soil-borne pathogens^30^. Climate data were taken from version 1.4 of the WorldClim dataset, which includes nineteen bioclimatic variables characterizing average and seasonal trends in temperature and precipitation^38^. In addition to climate data, we included layers describing elevation, soil, and two vegetation indices. Elevational data was taken from the Global Multi-resolution Terrain Elevation Data (GMTED2010) dataset provided by the US Geological Service. Soil layers were taken from the Global Soil Information Facilities (GSIF) SoilGrids database at 250 m resolution; four layers were included: soil pH, and the predicted soil content of sand, humult, and calcic vertisols at a depth of 0-5 cm^39^. The mean and amplitude of the normalized difference vegetation index (NDVI) were taken from the Trypanosomiasis & Land Use in Africa (TALA) dataset^40^. To prevent overfitting and reduce correlation among predictor variables, the variable set was reduced down using an automated procedure within the boosted regression tree implementation.

### Distribution Modelling

Boosted regression trees (BRTs) are currently considered a best practices method for modelling the global distribution of infectious diseases^11, 41^, including other soil-transmitted pathogens like *Burkholderia pseudomallei*^30^. In our study, BRTs were implemented using the ‘gbm’ package in R to develop a global species distribution model for *Bacillus anthracis*. Automated variable set reduction procedures selected a total of seventeen predictor layers: ten bioclimatic variables, two vegetation indices, elevation, and four soil variables. Presence, absence, and environmental data were run through the ‘gbm.step’ procedure on default settings (tree complexity = 4, learning rate = 0.005), following the established template of other studies. A total of 500 submodels were run; for each, presence points were bootstrapped, and pseudoabsences were randomly selected from the total 10,000 to achieve a 1:1 ratio^36^. Separate from the internal cross-validation (75%-25% split) of the BRT procedure, both presence and pseudoabsence points were split into an 80% training and 20% test dataset in each submodel, and model AUC was evaluated based on the independent test dataset. A final average model was calculated across the 500 submodels, and the 5^th^ and 95^th^ percentile were retained for use in the population at risk analyses. Models performed very well on the withheld test data (mean submodel AUC = 0.9244).

### Estimation of Human and Livestock Populations-at-Risk

We estimated the vulnerability of human and livestock populations by overlaying population datasets with maps of anthrax-suitable areas^42^. We mapped global anthrax suitability by dichotomizing model predictions with a threshold of 0.565 for suitable versus unsuitable predictions, with the threshold selected to maximize the true skill statistic (which weights sensitivity and specificity equally) of mean predictions across the entire dataset of all sightings and pseudoabsences. We estimated global population at risk using human population counts for 2015 from the Gridded Population of the World (GPW) dataset, version 4.0. We dichotomized “urban” and “rural” areas using the Global Human Built-up and Settlement Extent (HBASE) Dataset, version 1.0. We further split “urban” areas into urban and peri-urban based on the GPW population dataset, where we used a density under 1,000 persons per km^2^ as the threshold for classification as peri-urban. To measure possible occupational exposure, we used a global dataset of rural poor livestock keeper density.^43^ Finally, we estimated the number of livestock (cattle, sheep, goats, and swine) using a database of global livestock density at a spatial resolution of ∼1km x 1km (http://www.livestock.geo-wiki.org/).^44^ Those livestock populations at risk were further stratified by each of the livestock production zones using the livestock production systems data version 5 (http://www.livestock.geo-wiki.org/).^43–45^ For all human and livestock analyses, we obtained vulnerability estimates by overlaying population counts with dichotomized anthrax suitable areas from the BRT models; 95% credibility intervals were calculated by using the 5% (lower) and 95% (upper) bounds of the averaged BRT model prediction, in place of the mean prediction.^42^

### Delineating Wildlife at Risk

We identified ungulate species of interest based on species of conservation concern with known range overlap with anthrax, and measured the degree of range overlap with our global *B. anthracis* model, as a proxy for anthrax exposure. We selected seven candidate species of interest: pronghorn (*Antilocapra americana*), roan (*Hippotragus equinus*), saiga (*Saiga tatarica*), moose (*Alces alces*), reindeer (*Rangifer tarandus*), wild yak (*Bos mutus*), and bison (*Bison bison*). Of the seven, saiga are most threatened, and are listed on the IUCN red list as critically endangered. Yak are listed as vulnerable; bison are listed as near threatened; and pronghorn, roan, and moose are listed as least concern. We use the Global Protected Areas Database and official IUCN range maps, though we note that these range maps tend to overestimate ranges and can be misleading for conservation work^46^. (At least for saiga, we note that telemetry studies are currently underway to reassess the boundaries of the species’ range.)

## Acknowledgements

The authors thank: D. Pigott for helpful tips on boosted regression tree modelling; A. Barner for help obtaining WAHIS vaccination data; S.J. Ryan, for general feedback and technical support; G. Simpson, for visualization advice; P. Thornton, for access to the global dataset of rural poor livestock keepers; and T.A. Joyner, for data support; countless livestock and wildlife managers, clinicians, and field technicians for contributing data points. Partial funding for this study was provided by NIH 1R01GM117617-01 to JKB and WMG. This work was supported by the National Socio-Environmental Synthesis Center (SESYNC) under funding received from the National Science Foundation DBI-1639145.

## Author Contributions

CJC, ITK and JKB conceived of the study. JKB, MEHJ, ITK, and CJC collected and georeferenced data. CJC, ITK and JKB designed the models, and CJC ran models and analyses. NR contributed R code. All authors contributed to the writing and editing of the draft, and approved the study before submission.

